# Phase-field Approach to Cellular Blebbing

**DOI:** 10.1101/2025.03.14.643157

**Authors:** Kaihua Ji, Herbert Levine, Alain Karma

## Abstract

Bulges in the plasma membrane of cells known as blebs can form spontaneously in a wide range of biological processes but what controls their shape and stability remains incompletely understood. To address this, we introduce a dual phase-field model with coupled order parameters representing the cell cortex and plasma membrane that can quantitatively model blebbing in three dimensions. Simulations and analysis of the model reveal that, depending on whether blebbing occurs by detachment of the plasma membrane or rupture of the actin cortex, blebs can form discontinuously through a saddle node bifurcation or continuously with increasing cortical tension. The model predictions are in good quantitative agreement with existing experimental data for laser-induced cortex rupture.

---

Cellular blebbing is a process observed in a wide range of biological contexts from cell migration [1–3] to apoptosis [4], and has been extensively studied both experimentally and theoretically [5–8]. A key aspect of the blebbing phenomenology relates to disturbances of the actin cortex [9], which is a peripheral region of submicron thickness enclosing the cytogel (the bulk cytoskeleton and interstitial fluid) filling the volume of the cell. The cortex consists of an actomyosin network that generates an active tension *T* that is typically several fold larger than the passive membrane surface tension *γ*_*m*_.

Under resting conditions, the cortex is attached to the plasma membrane by specific anchoring proteins [10]. In contrast, in response to various perturbations the membrane can detach locally from the cortex, either by breaking of the anchoring bonds or more drastically by localized cortex failure. In these cases, the membrane will bulge outward to form a bleb. In the simplest conceptual scenario where the elasticity of the cortex and cytogel are both neglected, blebbing is predicted to be favored when the internal fluid pressure inside the cell ≈ 2*T/r*_*c*_ generated by the cortical tension, where *r*_*c*_ is the radius of a cell assumed to have a spherical shape, exceeds the Laplace pressure 2*γ*_*m*_*/r*_*b*_ of a hemispherical bleb protrusion of radius *r*_*b*_ *< r*_*c*_. Equating these two pressures yields the prediction that blebs form when *T* exceeds a critical value *T*_*c*_ ≈ *γ*_*m*_*r*_*c*_*/r*_*b*_. Taking into account elasticity slightly reduces the theoretical estimate of *T*_*c*_ [7], since part of the active tension is used up to compress the cortex and cytogel thereby producing a smaller increase of internal pressure, but does not change this basic picture.

While this current theory identifies several of the main biophysical parameters controlling bleb formation, it leaves some basic questions unanswered. Firstly, and perhaps most importantly, it does not explain in general what determines the value of *r*_*b*_. In the case of a *permanent* breakdown of the cortex in a circular region of radius *r*_*h*_, the natural assumption is that *r*_*b*_ ≈ *r*_*h*_ at *T* = *T*_*c*_ with further increase of bleb volume for *T > T*_*c*_. This makes sense for laser ablation experiments [7] which exogenously rupture the cortex, but this approach cannot explain the size of blebs produced endogenously, seemingly by transient disturbances in the cortex-membrane interaction. Even the question of whether such an endogenous bleb can continue to exist after such a transient event and if so, over what range of tension, has remained unanswered. Secondly, it remains unclear how the membrane-cortex attachment strength affects the answer to this question since this important biophysical parameter does not enter in the current theory of exogenous blebbing. To address these questions, we develop a phase-field (PF) approach [11] that has proven successful in modeling complex interfacial patterns in a wide range of physical [12–14] and biological [15] contexts. We describe the cell membrane and cortex by scalar fields *ϕ*_*m*_ and *ϕ*_*c*_, respectively, which vary smoothly from 0 outside the cell or cortex to 1 inside. We incorporate the forces acting on the cell and cortex by adding separate contributions to a functional representing the total energy *U* of the cortex-membrane-cytogel system

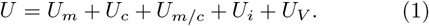

The individual contributions

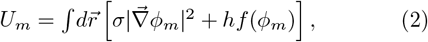

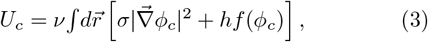

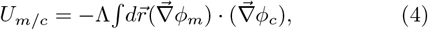

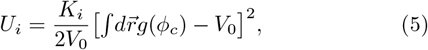

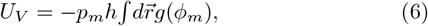

represent the passive membrane energy (*U*_*m*_), the energy associated with active cortex tension (*U*_*c*_), the binding energy of the membrane to the cortex (*U*_*m/c*_), the elastic energy of the cytogel (*U*_*i*_) where *K*_*i*_ is the bulk modulus that includes the contribution of both the cytogel and cortex elasticity [16] and *V*_0_ a reference equilibrium volume; *U*_*V*_ constrains the cell volume 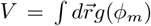 to remain constant through the Lagrange multiplier *p*_*m*_. In addition, *f* (*ϕ*) = 4*ϕ*^2^(1 ™ *ϕ*)^2^ is a standard double-well potential with minima at *ϕ* = 0 and *ϕ* = 1, and *g*(*ϕ*) = *ϕ*^3^[10 + 3*ϕ*(2*ϕ ™* 5)] is a nonlinear function that increases monotonically with *ϕ* and has vanishing first and second derivatives at *ϕ* = 0 and *ϕ* = 1. The PF evolution equations have the variational from

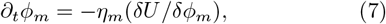

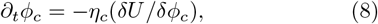

which drives the coupled dynamics of the membrane and cortex towards a global minimum of *U* under the constant volume constraint 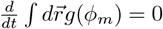 together with Eq. (7), this constraint uniquely determines *p*_*m*_ representing the spatially uniform internal pressure required to maintain the cell volume constant.

To relate the PF model and physical parameters, we use standard expressions for the excess energies of locally planar interfaces to obtain the following expressions for the passive membrane tension *γ*_*m*_ = 2 ⎰ *σ* (∂_*x*_ *ϕ* _*m*_)^2^ *dx*, the active cortex tension *γ*_*c*_ = 2 ⎰ *νσ* (∂_*x*_*ϕ*_*m*_)^2^ *dx*, and the combined tension of the membrane and cortex bound to each other

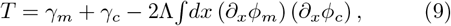

where *ϕ*_*m*_(*x*) and *ϕ*_*c*_(*x*) are the stationary one-dimensional PF profiles along a direction *x* normal to these interfaces. When the membrane and cortex are detached (*U*_*m/c*_ = 0), both *ϕ*_*m*_(*x*) and *ϕ*_*c*_(*x*) reduce to the standard tangent hyperbolic profile *ϕ*_0_(*x*) = [1 − tanh(*x/W*)]*/*2, where *W* ≡ (*σ/h*)^1*/*2^ is the interface thickness, yielding 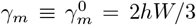 and 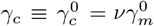 [16]. When the cortex and membrane are colocalized, the contribution of the binding energy *U*_*m/c*_ slightly modifies the stationary PF profiles. However, they remain well approximated by *ϕ*_0_(*x*) and an expression for *T* is obtained by substituting *ϕ*_*m*_(*x*) = *ϕ*_*c*_(*x*) = *ϕ*_0_(*x*) into Eq. (9), which yields

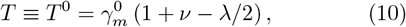

where we have defined the dimensionless membrane-cortex bonding strength *λ ≡* Λ*/*(*hW* ^2^). Since the values of *γ*_*m*_, *γ*_*c*_, and *T* differ from 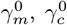, and *T* ^0^ only by a small amount that increases with *λ* when the cortex and membrane are bound [16], the analytical expressions with superscript ^0^ can be used to relate model and physical parameters. For our simulations, we first use generic physical parameters with a cell radius *r*_0_ = 8 *µ*m, *K*_*i*_ = 750 Pa, and 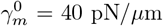. The interface thickness *W* is the only free parameter of the model. Once *W* is chosen, the barrier height *h* of the double-well potential can always be adjusted to match a desired physical value of 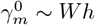∼ *Wh*, and thus 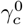 and *T* ^0^ by further choosing the dimensionless parameters 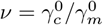 and *λ* using Eq. (10). *W* is typically chosen as large as possible to keep the computations tractable, but small enough for the results to be independent of *W* . Satisfactory convergence was obtained with *W* = 0.213 *µ*m that is used in all simulations unless explicitly stated. Finally, we only use here the evolution Eqs. (7)-(8) to model stationary blebs corresponding to global minima of *U* where all forces exactly balance each other, allowing the kinetic parameters *η*_*m*_ and *η*_*c*_ to be arbitrarily set around unity.

We first investigate bleb formation without cortex rupture. To produce blebs, we enable the membrane to detach transiently from the cortex by setting *λ* = 0 over a small circular heterogeneous region of radius *r*_*h*_ on the periphery of the cell. Furthermore, we choose *ν* large enough so that the tension *T* is above the threshold 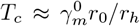 where the internal pressure exceeds the Laplace pressure of a hemispherical bleb. Results of simulations for *ν* = 16 and *r*_*h*_*/r*_0_ = 0.08 and different *λ* values are shown in Fig. 1. Fig. 1(a) illustrates how the bleb morphologies change with *λ* and Fig. 1(b) illustrates the PF profiles for one morphology. Importantly, these blebs remain stationary even after the transient detachment perturbation that enabled their formation is switched off by resetting *λ* inside the disk region of radius *r*_*h*_ to the same finite value used outside this region (equivalent to setting *r*_*h*_ = 0). Hence, these *endogenous* blebs exist even when the entire cortex is still intact and able to anchor the membrane. This is because their morphologies are controlled by the balance of forces at the circular contact line between the the bleb and cortex. As for a liquid droplet on a surface, the wetting behavior of the bleb is controlled by Young’s condition *T* = *γ*_*m*_ cos *θ* + *γ*_*c*_ which, combined with Eq. (10) and the approximations 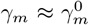 and 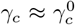, yields at once the prediction for the contact angle *θ* of the bleb on the cortex

**FIG. 1.**
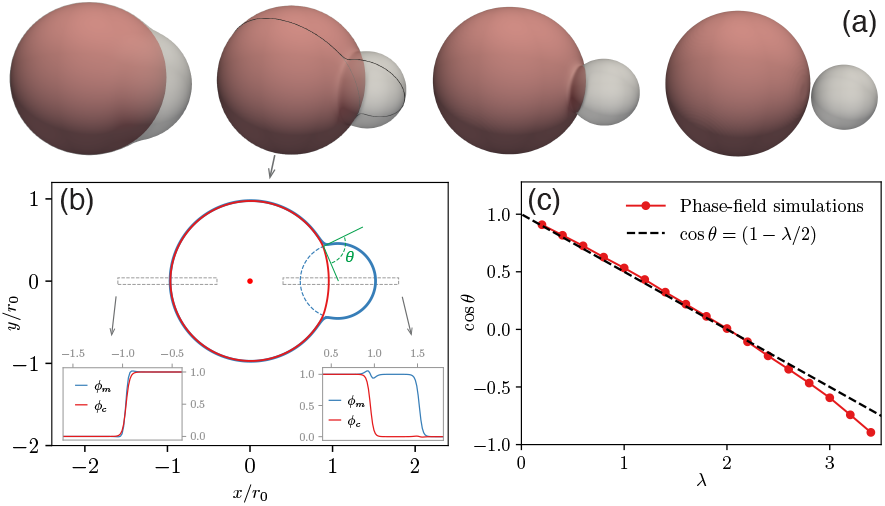
(a) Simulated 3D endogenous bleb morphologies that remain stable after a transient suppression of membrane-cortex attachment in a small circular region of radius *r*_*h*_*/r*_0_ = 0.08 for *ν* = 16 and *λ* = 0.5, 2, 3, and 3.5 increasing from left to right. The blebs remain stationary after attachment is restored on the whole cortex (*r*_*h→*_ 0). (b) Membrane (*ϕ*_*m*_) and cortex (*ϕ*_*c*_) phase-field (PF) profiles in a 2D cross-section of the 3D bleb morphology for *λ* = 2. (c) Comparison of contact angle *θ* extracted from simulations and analytically predicted.

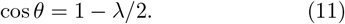

Fig. 1(c) shows that the contact angles extracted from the simulated blebs are in good quantitative agreement with the prediction of Eq. (11). The discrepancy at larger value of *λ* is due to the departure of the PF profiles from *ϕ*_0_(*x*) [Fig. 1(b)], which causes tensions to deviate from their analytical values.

To investigate in more detail the nature of the nonlinear bifurcation to endogenous blebs with intact cortex, we carry out a series of simulations, initially for *λ* = 2. Here, we again start with a small hole of radius *r*_*h*_ and slowly increase the tension–the data is presented as blue filled triangles in Fig. 2(a)-(b), showing respectively the bleb radius of curvature and volume normalized by the initial cell radius *r*_0_ and volume 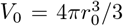. Up to a critical tension close to *T*_*c*_, blebs remain constrained to the hole with a bleb radius decreasing from *r*_*b*_ ≈ *r*_0_ for low tension to *r*_*b*_ ≈ *r*_*h*_ at the critical tension. Just past the critical tension, the bleb size (measured by both *r*_*b*_ and *ρ*) has a discontinuous jump resulting from the fact that the bleb contact line with the cortex is no longer constrained to the border of the detached circular region. Instead, the perimeter of the contact line grows by further detaching the membrane from the cortex over a larger region until force balance set by Young’s condition is satisfied. At this point, the hole can be eliminated, leaving an endogenous bleb. Thus, to investigate this upper branch of endogenous bleb solutions, we switch off the perturbation (i.e., set *r*_*h*_ = 0), and both increase and decrease the tension. The results shown as open red circles in Fig. 2(a)-(b) reveal that endogenous blebs continue to exist for significantly lower tension than *T*_*c*_ and vanish discontinuously when *T* falls below a lower critical tension 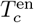. For 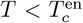, the membrane reattaches to the cortex corresponding to red circles with *r*_*b*_ = *r*_0_ and *ρ* = 0 in Fig. 2(a)-(b). Repeating the series of simulations for different values of *λ*, we obtain 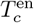 as a function of contact angle *θ* as shown in Fig. 2(c).

**FIG. 2.**
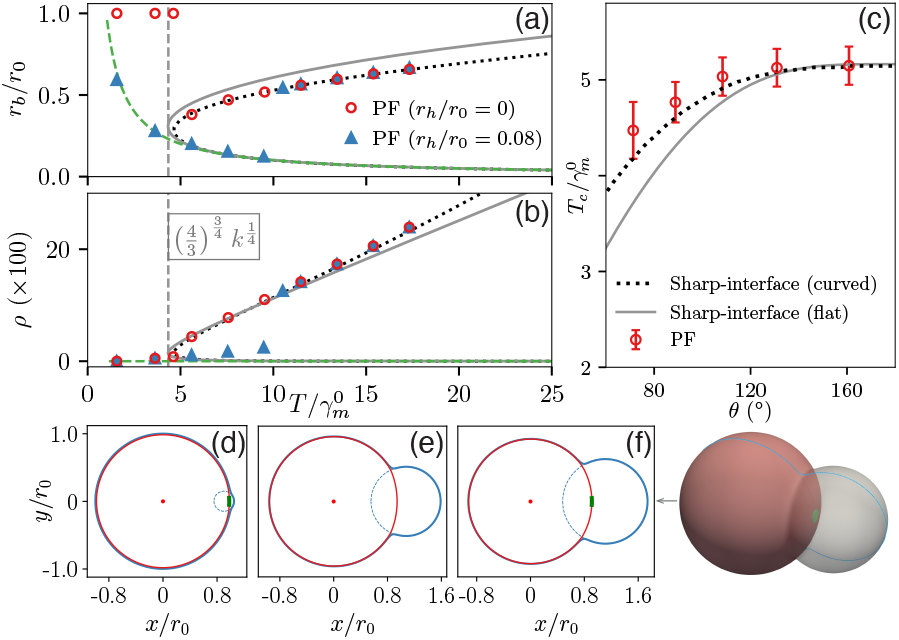
(a) Scaled bleb radius *r*_*b*_*/r*_0_ and volume *ρ* = *V*_*b*_*/V*_0_ (b) versus scaled membrane-cortex tension 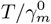 for *λ* = 2. Symbols correspond to 3D PF simulations and the lines to analytical predictions without (solid gray line) and with (dotted lines computed numerically [16]) cortex curvature; the green dashed line corresponds to 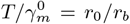 . (c) 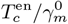 versus contact angle *θ* from PF simulations and analytical predictions. 2D cross-sections of PF profiles illustrating that a small bleb (d) and a large bleb (e) can co-exist at the same tension 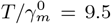 in the presence (d) and absence (e) of a circular region (*r*_*h*_*/r*_0_ = 0.08) with suppressed membrane-cortex binding marked as a green patch. (f) 2D cross-section for a large tension 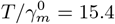.

To obtain an analytical understanding of the bifurcation to endogenous blebs, we consider the sharp-interface (*W/r*_0_ → 0) limit of the PF model. In this limit, the energy *U* of the membrane-cortex-cytogel system (Eq. 1) reduces to

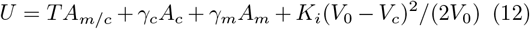

where *A*_*m/c*_ and *A*_*c*_ are the areas of the membrane-bound and unbound cortex regions, respectively, *A*_*m*_ is the area of the detached membrane, and 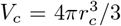 is the volume of the cytogel inside the cortex. Minimization of Eq. (12) with respect to the bleb radius *r*_*b*_ under the constraints of (i) total volume conservation *V* = *V*_*c*_+*V*_*b*_ = *V*_0_, and (ii) Young’s condition *T* = *γ*_*m*_ cos *θ* + *γ*_*c*_ at the bleb/cortex contact line, uniquely fixes *U* as a function of *r*_*b*_. Minimization of *U* with respect to *r*_*b*_, i.e., *dU/dr*_*b*_ = 0, yields a relation between bleb radius and tension [16]

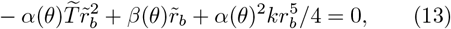

where we have defined the functions *α*(*θ*) = 1−3 cos *θ/*2+ cos^3^ *θ/*2 and *β*(*θ*) = 1 − cos *θ*(1 +sin^2^ *θ/*2), and the scaled quantities 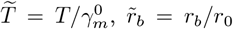, and 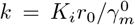. Eq. (13) predicts that endogenous blebs emerge through a saddle node bifurcation where an upper stable and lower unstable branch of stationary solutions [solid lines in Fig. 2(a)-(b)] meet at a lower critical tension. The latter is determined by the dual condition 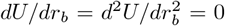, which yields the prediction for the saddle node bifurcation point [16]

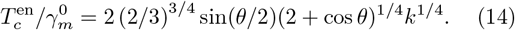

This prediction is plotted as a solid line in Fig. 2(c). The modest quantitative discrepancy between analytical and PF results stems from the fact that the expression for *U* used to derive Eqs. (13)-(14) assumes that the cortex is locally planar, as opposed to spherical, in the region of the bleb contact line, which is only asymptotically valid in the limit *r*_*b*_*/r*_0_ → 0. A more elaborate geometrical calculation that takes into account cortex curvature [16] (requiring *r*_*b*_ to be solved numerically) yields the dashed lines in Fig. 2(a)-(c), which are in excellent agreement with PF results.

Next, we further consider *exogenous* blebs forming in the presence of a permanent (i.e., persisting on the time scale of bleb formation) perturbation that causes local membrane detachment in a small region of cortex. We choose *λ* = 3.5 such that outside this region, strong membrane-cortex binding constrains the bleb contact line to the periphery of the detached region; note that the contact angle is now *π* (see Fig. 1c), leaving the only endogenous solution as expulsion of the entire bleb from the original cell. This has the desired effect for the exogenous bleb of constraining the contact line to have a radius close to *r*_*h*_ over the range of *T* where the bifurcation occurs. As before, we increase the tension until the bleb volume makes a sudden jump and then both increase and decrease the tension to look for hysteresis. Simulation results for exogenous blebs in Fig. 3 show a hysteretic bifurcation picture with a sudden jump in bleb volume when *T* exceeds 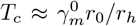, and with blebs persisting to a lower tension 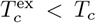 when *T* is decreased. The bifurcation can again be analyzed in the sharp-interface limit by minimization of the total energy *U* with the bleb-cortex contact line now anchored at the periphery of the detached region with a fixed radius *r*_*h*_, instead of moving freely on the cortex to satisfy Young’s condition (11) as for endogenous blebs. The results of this calculation yield the prediction [16]

**FIG. 3.**
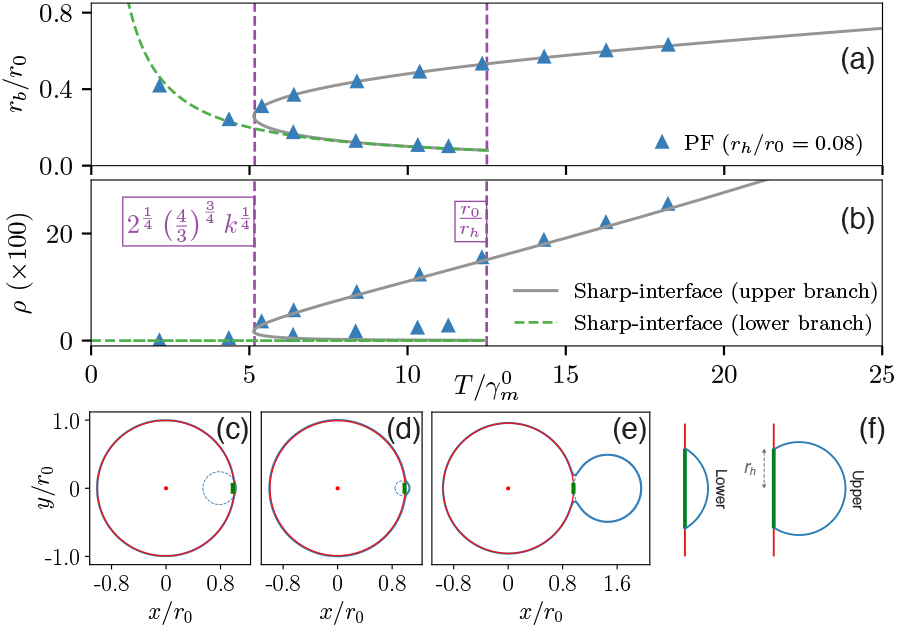
Similar bifurcation diagram as Fig. 2 but for a larger value *λ* = 3.5 where the bleb contact line remains confined to the periphery of the circular region with a permanent suppression of membrane-cortex binding. (a) *r*_*b*_*/r*_0_ and *ρ* = *V*_*b*_*/V*_0_ (b) versus 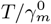. (c)-(e) cross-sections for 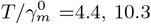, and 10.3, respectively, where the small and large blebs shown in (d) and (e) coexist at the same tension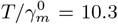. (f) Schematics of bleb profiles corresponding to the lower and upper branches of stationary solutions.

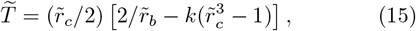

with

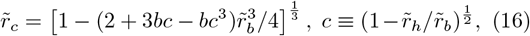

where all radii are scaled by *r*_0_ as before. Eqs. (15)-(16) uniquely determine the bleb radius 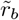 as a function of tension 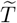 with *b* = ™1 for the lower stable branch and *b* = +1 for the middle unstable and upper branches of stationary solutions. Fig. 3(a)-(b) show that the predicted lower (green dashed line) and upper (black solid line) branches of solutions are in excellent quantitative agreement with PF results. In the limit *r*_*h*_*/r*_0_ → 0, the lower critical tension at which the upper stable and middle unstable branches of exogenous blebs meet, 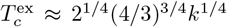, coincides with the *θ → π* limit of the endogenous bleb prediction 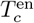 given by Eq. (14), consistent with the fact that the bleb-cortex contact line shrinks to zero radius in both limits.

Finally, we consider the case where membrane detachment is triggered by cortex rupture over a hole region of radius *r*_*h*_ and use physical parameters corresponding to the laser ablation experiments [7] (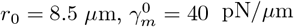 and *K*_*i*_ = 2283 Pa). Since *ϕ*_*c*_ is a continuous field, the present PF approach cannot model explicitly a hole in the cortex. However, as long as detachment occurs over a circular region, the blebbing bifurcation is expected to be quantitatively the same whether this region is a cortex unable to bind the membrane or an actual hole as far as stationary blebs determined by force balance are concerned. The main difference is that the hole expands significantly with increasing tension. Interestingly, results in Fig. 4 show that this expansion, which has been experimentally observed and theoretically predicted from elastic shell theory [7], can affect the blebbing bifurcation. To illustrate this, we first show in Fig. 4(a)-(b) the predictions of Eqs. (15)-(16) for different fixed values of *r*_*h*_. The blebbing bifurcation as a function of tension is seen to change from being discontinuous and hysteretic for smaller *r*_*h*_ to continuous for larger *r*_*h*_ in agreement with PF simulations. This change of bifurcation character stems from the fact that 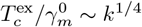is determined predominantly by the cytogel compressibility, but is independent of *r*_*h*_, while 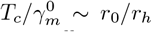 is predominantly determined by *r*_*h*_ up to a small correction of order 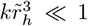 [16]. Therefore, as *r*_*h*_ increases, *T*_*c*_ decreases and the transition becomes continuous when 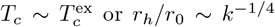 up to a prefactor of order unity.

**FIG. 4.**
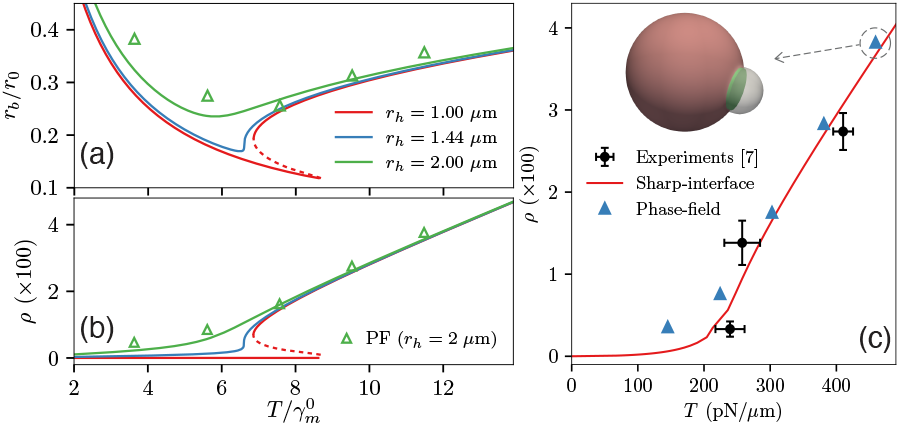
Analytical predictions of Eqs. (15)-(16) for *r*_*b*_*/r*_0_ and *ρ* (b) illustrating how, when *r*_*h*_ is increased, the blebbing transition changes from discontinuous and hysteretic (red solid and dashed lines) to continuous and non-hysteretic (green line). Symbols represent PF simulations with *λ* = 2 and *W* = 0.16 *µ*m. (c) Analytical predictions of Eqs. (15)-(16) and PF simulations using the measured linear relationship 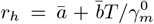 in the laser ablation experiments [7] 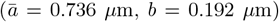 and corresponding measurements of *ρ* in the same experiments.

As a last step to compare quantitatively our predictions to experiments, we use as input into the model the measured linear dependence of hole radius with tension from [7]. Fig. 4(c) shows that the results of both PF simulations and sharp-interface theory [Eqs. (15)-(16)] are in remarkably good quantitative agreement with experiments with no adjustable parameter. Interestingly, the hole expansion renders the bifurcation continuous for this particular set of experimental parameters. However, it is clear from the above predictions, which should be experimentally testable, that the bifurcation could be hysteretic for other parameters such as a smaller hole size or lower cytogel compressibility. The present PF approach can also be applied to model the formation of multiple blebs. Simulation results show that configurations with two endogenous or exogenous blebs from the upper branches of stationary solutions in Figs. 2-3 are unstable and evolve towards a single bleb of larger volume, while small exogenous blebs from the lower branch of solution (e.g., Fig. 3c or d) can be stable, consistent with analytical predictions based on energy minimization [16].

The present results demonstrate the power of the PF approach to model complex 3D shape changes associated with local detachment of the membrane from the cortex under the combined effects of active and passive forces. While we have focused here on static aspects of blebbing, the approach could be extended to investigate important dynamical aspects such as the role of blebs in amoeboid cell motility [17]. This should be possible by extending the present framework to incorporate fluid flow [8, 18], reassembly of the contractile actin cortex in blebs [19], and cell-membrane-substrate interaction [20]. Work along this line is currently in progress.

This work was supported by the National Science Foundation (NSF) under grant PHY-2412650; additional support was obtained from the Center for Theoretical Biological Physics, NSF PHY-2019745. K.J. acknowledges partial support from Lawrence Livermore National Laboratory (LLNL) under contract DE-AC52-07NA27344. Numerical simulations were performed on the Discovery cluster of Northeastern University located in Massachusetts Green High Performance Computing Center (MGHPCC) in Holyoke, MA.

## Supporting information

The Supplemental Material includes additional simulations and analyses.

