## Supplementary material for "Phase-field Approach to Cellular Blebbing": The Supplemental Material includes additional simulations and analyses.

#### CONTENTS

I. Dual phase-field model for cellular blebbing 1
  A. Evolution equations 1
  B. Stationary solutions 2
  C. Interfacial free-energy 3
  D. Scaling of the model 4
  E. Numerical implementation 5

II. Sharp-interface theory 6
  A. Volume conservation 6
  B. Endogenous bleb 7
    1. Equilibrium solutions 7
    2. Critical tensions 7
  C. Exogenous bleb 9
    1. A simple picture 9
    2. $r_h \to 0$ limit 9
    3. Arbitrary contact angle 10

III. Multiple blebs 11
  A. Phase-field simulations of two blebs 11
  B. Stability of two-bleb configurations 11
    1. Endogenous blebs 11
    2. Exogenous blebs 12

References 13

#### I. DUAL PHASE-FIELD MODEL FOR CELLULAR BLEBBING

In this supplementary section, we first present the derivation of evolution equations of a dual phase-field (PF) model. Subsequently, we derive the stationary solutions and the interfacial free-energy. Then, we show the scaling of the model and the numerical implementation.

##### A. Evolution equations

We define a PF, $\phi_m$, to represent the cell membrane, which varies smoothly from $\phi_m = 1$ within the cell interior to $\phi_m = 0$ outside. The passive membrane energy is expressed as:

$$U_m = \int d\vec{r} \left[\sigma|\vec{\nabla}\phi_m|^2 + hf(\phi_m)\right] . \tag{S1}$$

Inside the integrand, the first term is the gradient energy that ensures a finite interface thickness, and the second term is a double-well potential providing the stability of the two phases ($\phi_m = 0$ and 1), where $h$ is the energy barrier height and $f(\phi) = 4\phi^2(1-\phi)^2$. Analogously, the cell cortex is represented by another phase field, $\phi_c$, which takes on a value of $\phi_c = 1$ within the cortex, and $\phi_c = 0$ outside. The energy corresponding to the cell cortex is then given by:

$$U_c = \nu \int d\vec{r} \left[\sigma|\vec{\nabla}\phi_c|^2 + hf(\phi_c)\right], \tag{S2}$$

where $\nu$ is a dimensionless coefficient for controlling the cortical active tension. We further consider the energy contribution due to membrane-cortex interaction:

$$U_{m/c} = -\Lambda \int d\vec{r}(\vec{\nabla}\phi_m) \cdot (\vec{\nabla}\phi_c), \tag{S3}$$

where $\Lambda$ is a coupling coefficient. Lastly, the elastic energy arising from the compressible cytogel is given by:

$$U_i = \frac{K_i}{2V_0}\left(\int d\vec{r} g(\phi_c) - V_0\right)^2, \tag{S4}$$

where $K_i$ denotes the bulk modulus of the cytogel, $g(\phi) = \phi^3[10 + 3\phi(2\phi - 5)]$, and $V_0$ is the initial cell volume.

The total energy of this membrane-cortex-cytogel system is

$$U = U_m + U_c + U_{m/c} + U_i + U_V, \tag{S5}$$

representing the contributions from membrane, cortex, the cortex-membrane interaction, the elastic cytogel, and an additional contribution

$$U_V \equiv -p_m h \int d\vec{r} g(\phi_m) \tag{S6}$$

used to maintain the cell volume (the volume confined by membrane) a constant, where $p_m$ is a Lagrange multiplier. The evolution equations are derived from standard variational dynamics

$$\frac{\partial\phi_m}{\partial t} = -\eta_m \frac{\delta U}{\delta\phi_m}, \tag{S7}$$

and

$$\frac{\partial\phi_c}{\partial t} = -\eta_c \frac{\delta U}{\delta\phi_c}. \tag{S8}$$

As a result, we obtain the evolution equations for both phase fields $\phi_m$ and $\phi_c$:

$$\frac{1}{\eta_m}\frac{\partial\phi_m}{\partial t} = 2\sigma\nabla^2\phi_m - hf'(\phi_m) - \vec{\nabla}\cdot\left(\Lambda\vec{\nabla}\phi_c\right) + hp_m g'(\phi_m), \tag{S9}$$

and

$$\frac{1}{\eta_c}\frac{\partial\phi_c}{\partial t} = 2\nu\sigma\nabla^2\phi_c - \nu hf'(\phi_c) - \vec{\nabla}\cdot\left(\Lambda\vec{\nabla}\phi_m\right) - \frac{K_i}{V_0}\left(\int d\vec{r} g(\phi_c) - V_0\right) g'(\phi_c), \tag{S10}$$

where the coupling coefficient $\Lambda$ can be either a constant or a spatially varying scalar field. The Lagrange multiplier $p_m$ is used to maintain the cell volume and is calculated through the relation $\frac{d}{dt}\int d\vec{r} g(\phi_m) = 0$.

##### B. Stationary solutions

For an isolated phase field $\phi_m(x)$ without the membrane-cortex interaction in 1D, the equilibrium condition is

$$\frac{\partial\phi_m}{\partial t} = 0 = 2\sigma\partial_x^2\phi_m - hf'(\phi_m). \tag{S11}$$

With the boundary conditions $\phi_m(x = +\infty) = 0$ and $\phi_m(x = -\infty) = 1$, we obtain the stationary solution as a function of $x$:

$$\phi_m^0(x) = \frac{1}{2}\left[1 - \tanh\left(\frac{x - x_0}{W}\right)\right], \tag{S12}$$

where $W \equiv (\sigma/h)^{1/2}$ is the interface thickness, and $x_0$ is the location of membrane, i.e., the location where $\phi_m^0(x_0) = 1/2$. Similarly, one can obtain the stationary solution for the cortex phase field:

$$\phi_c^0(x) = \frac{1}{2}\left[1 - \tanh\left(\frac{x - x_0}{W}\right)\right]. \tag{S13}$$

##### C. Interfacial free-energy

When membrane and cortex are bound to each other, the combined tension $T$ includes contributions from membrane ($U_m$), cortex ($U_c$), and the membrane-cortex interaction ($U_{m/c}$). According to Eqs. (S1)-(S3), we obtain

$$T = \int dx \left[\sigma\left(\partial_x \phi_m\right)^2 + hf\left(\phi_m\right) + \nu\sigma\left(\partial_x \phi_c\right)^2 + \nu hf\left(\phi_c\right) - \Lambda_{m/c}\partial_x\phi_m\partial_x\phi_c\right], \tag{S14}$$

Here we have considered a 1D case and the $x$ direction is perpendicular to the interfaces. Under the assumption that the cytogel compression is negligible, the equilibrium conditions of membrane and cortex phase fields are:

$$2\sigma\partial_x^2\phi_m - hf'(\phi_m) - \Lambda_{m/c}\partial_x^2\phi_c = 0, \tag{S15}$$

and

$$2\nu\sigma\partial_x^2\phi_c - \nu hf'(\phi_c) - \Lambda_{m/c}\partial_x^2\phi_m = 0. \tag{S16}$$

Let Eq. (S15) multiplied by $\partial_x\phi_m$ and Eq. (S16) multiplied by $\partial_x\phi_c$, and we obtain

$$\partial_x\left[\sigma\left(\partial_x\phi_m\right)^2 - hf\left(\phi_m\right)\right] - \Lambda_{m/c}\partial_x\phi_m\partial_x^2\phi_c = 0, \tag{S17}$$

and

$$\nu\partial_x\left[\sigma\left(\partial_x\phi_c\right)^2 - hf\left(\phi_c\right)\right] - \Lambda_{m/c}\partial_x\phi_c\partial_x^2\phi_m = 0. \tag{S18}$$

Summing up both sides of Eqs. (S17)-(S18) yields

$$\partial_x\left[\sigma\left(\partial_x\phi_m\right)^2 - hf\left(\phi_m\right) + \nu\sigma\left(\partial_x\phi_c\right)^2 - \nu hf\left(\phi_c\right) - \Lambda\partial_x\phi_m\partial_x\phi_c\right] = 0 \tag{S19}$$

With conditions $\partial_x\phi_m = \partial_x\phi_c = f(\phi_m) = f(\phi_c) = 0$ in bulk phases ($\phi_m$ or $\phi_c = 0$ or 1), we obtain

$$hf\left(\phi_m\right) + \nu hf\left(\phi_c\right) = \sigma\left(\partial_x\phi_m\right)^2 + \nu\sigma\left(\partial_x\phi_c\right)^2 - \Lambda\partial_x\phi_m\partial_x\phi_c. \tag{S20}$$

Combining Eq. (S14) and (S20), we derive a general expression for the surface tension:

$$T = 2\int dx\left[\sigma\left(\partial_x\phi_m\right)^2 + \nu\sigma\left(\partial_x\phi_c\right)^2 - \Lambda\left(\partial_x\phi_m\right)\left(\partial_x\phi_c\right)\right]. \tag{S21}$$

Specifically, within regions where $\phi_c = 0$, the tension is $\gamma_m = hWa_1$, where $a_1 \equiv 2W\int dx\left[(\partial_x\phi_m)^2\right]$ is a dimensionless coefficient. With the stationary solution $\phi_m^0(x)$ in Eq. (S12), we obtain $a_1 \equiv a_1^0 = 2/3$ and $\gamma_m \equiv \gamma_m^0 = hWa_1^0$. Conversely, for regions where $\phi_m = 0$, the tension is $\gamma_c \equiv \gamma_c^0 = \nu hWa_1^0$. In other regions, $T \equiv T^0 = \gamma_m^0(1 + \nu - \lambda/2)$. As shown in Fig. S1, the values of $\gamma_m$, $\gamma_c$, and $T$ differ from $\gamma_m^0$, $\gamma_c^0$, and $T^0$ only by a small amount that increases with $\lambda$ when the cortex and membrane are bound.

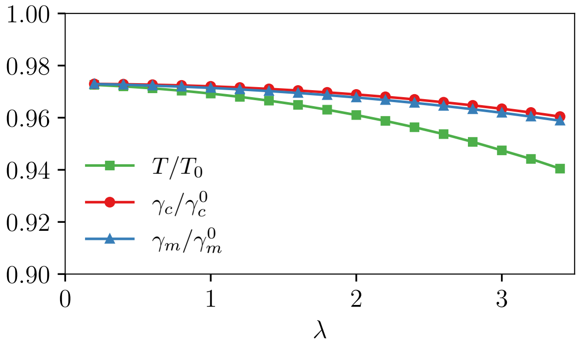

FIG. S1. The ratio between the measured tensions in 3D PF simulations with different $\lambda$ and their stationary solutions. Simulations are performed for $\nu = 16$, corresponding to Fig. 1(c) in the main text.

##### D. Scaling of the model

To facilitate the numerical implementation, we scale evolution equations (S9)-(S10) to dimensionless forms:

$$\frac{\partial \phi_m}{\partial t} = 2\nabla^2\phi_m - f'(\phi_m) - \vec{\nabla}\cdot\left(\lambda\vec{\nabla}\phi_c\right) + p_m g'(\phi_m), \tag{S22}$$

and

$$\alpha^{-1}\frac{\partial \phi_c}{\partial t} = 2\nu\nabla^2\phi_c - \nu f'(\phi_c) - \vec{\nabla}\cdot\left(\lambda\vec{\nabla}\phi_m\right) - \frac{k_i}{v_0}\left(\int d\vec{r} g(\phi_c) - v_0\right) g'(\phi_c), \tag{S23}$$

where the time is scaled by $\tau_0 \equiv 1/(\eta_m h)$, and the length is scaled by the interface thickness, $W \equiv (\sigma/h)^{1/2}$. The dimensionless coefficients include: $\lambda \equiv \Lambda/(hW^2)$, $\alpha \equiv \eta_c/\eta_m$, $k_i \equiv K_i/h$, and $v_0 \equiv V_0/W^3$. The Lagrange multiplier $p_m$ is updated at each iteration according to the relation

$$\frac{d}{dt}\int d\vec{r} g(\phi_m) = \int d\vec{r} g'(\phi_m)\frac{\partial \phi_m}{\partial t} = 0, \tag{S24}$$

where $\partial_t\phi_m$ is substituted by the right-hand-side of Eq. (S22). The explicit equation for calculating $p_m$ is

$$p_m = \frac{-\int d\vec{r} g'(\phi_m)\left[2\nabla^2\phi_m - f'(\phi_m) - \vec{\nabla}\cdot\left(\lambda\vec{\nabla}\phi_c\right)\right]}{\int d\vec{r} g'(\phi_m) g'(\phi_m)}. \tag{S25}$$

Since the interface thickness $W$ is a free parameter in the PF model, we need to investigate how dimensionless coefficients $k$ and $\lambda$ scale with $W$. In Section I C, we showed that the membrane free-energy is $\gamma_m^0 = hWa_1^0$ with the stationary PF profile. Combining with the relation $K_i = k_i h$, we can eliminate $h$ and obtain

$$k_i = a_1^0\frac{WK_i}{\gamma_m^0}. \tag{S26}$$

Thus, for given physical parameters $K_i$ and $\gamma_m^0$, the coefficient $k_i$ varies proportional to $W$. With stationary solutions $\phi_m^0(x)$ and $\phi_c^0(x)$ in Eqs. (S12)-(S13), we compute the membrane-cortex interaction energy to be

$$U_{m/c} = -\Lambda\int dx(\partial_x\phi_m^0)\cdot(\partial_x\phi_c^0) = \frac{1}{3}\lambda hW^3. \tag{S27}$$

Combing $\gamma_m^0 = hWa_1^0$ and Eq. (S27), we eliminate $h$ and obtain $U_{m/c} = -\lambda W^2\gamma_m^0/2$. Thus, for a given value of $\gamma_m^0$, $U_{m/c} \sim \lambda W^2$. Similarly, one can derive that $U_m \sim W^2$ and $U_c \sim W^2$. In the PF simulation, it is the relative ratio of $U_m$, $U_c$, and $U_{m/c}$ (or equivalently the Young's condition $T = \gamma_c + \gamma_m\cos\theta$) that determines the contact angle $\theta$. Thus, $\theta$ is independent of $W$, and it is uniquely determined by $\lambda$.

In Table S1, we listed the physical and modeling parameters that correspond to three values of interface thickness: $W = 0.16$ $\mu$m, $W = 0.213$ $\mu$m, and $W = 0.32$ $\mu$m. In simulations with different $W$, the changed parameters include: the initial cell radius $r_0$ (in the unit of $W$) that scales with $W^{-1}$, and $k$ that scales with $W$ according to Eq. (S26). In Fig. S2, we compare the shapes of cell membranes and cortex in PF simulations with different values of $W$, where the convergence of cell shapes is excellent. However, as we will show in Fig. S4, the PF simulation requires a small enough $W$ to resolve the critical tension $T_c$ for blebbing, especially for a contact angle $\theta < 90°$.

TABLE S1. Physical and modeling parameters.

| Symbol | Description | | Value | | Unit | Ref. |
|---|---|---|---|---|---|---|
| **Physical parameters** | | | | | | |
| $r_0$ | Cell radius | | 8 | | $\mu$m | [1] |
| $\gamma_m^0$ | Interfacial free-energy of membrane | | $4 \times 10^{-17}$ | | J $\mu$m$^{-2}$ | [2, 3] |
| $K_i$ | Bulk modulus | | $7.5 \times 10^{-16}$ | | J $\mu$m$^{-3}$ | |
| **Modeling parameters** | | | | | | |
| $\Delta x$ | Grid spacing | | 0.4 | | $W$ | |
| $\alpha$ | $\eta_c/\eta_m$ | | 0.8 | | | |
| $R_t$ | Coefficient for numerical stability | | 0.5 | | | |
| **Parameters scaled with** $W$ | | | | | | |
| $W$ | Interface thickness | 0.16 | 0.213 | 0.32 | $\mu$m | |
| $h$ | Energy barrier height | 375 | 281.25 | 187.5 | J m$^{-3}$ | |
| $r_0$ | Initial cell radius | 50 | 37.5 | 25 | $W$ | |
| $k_i$ | Scaled bulk modulus of cortex $K_i/h$ | 2 | 2.6667 | 4 | | |

Here, we discuss the value of bulk modulus $K_i$ in PF simulations. The compressibility of a cell has contributions from both cytogel and cortex. The force balance is described by a relation [2]:

$$\frac{2T}{r_c} - \left(\frac{E_i}{1-2v_i} + 4E_c h_c \frac{1}{r_c}\right)\frac{\Delta r_c}{r_c} = \frac{2\gamma_m}{r_b}, \quad \text{(S28)}$$

where $r_b$ is the bleb radius, $r_c$ is the cell radius, $\Delta r_c$ is the variation of the cell radius due to bleb formation, $E_i$ is the Young's modulus of the cytoplasm, $E_c$ is the Young's modulus of the actin cortical gel, $v_i$ is Poisson's ratio, and $h_c$ is the cortex thickness. Effectively, the term $\frac{E_i}{1-2v_i}$ represents the cytoplasmic elasticity, and $E_c h_c$ corresponds to the cortex elastic modulus. The contribution of the cortex is relatively small since $E_i$ and $E_c$ are comparable while $h_c/r_c \ll 1$. The total elasticity fitting experimental data for L929 cells is $\left(\frac{E_i}{1-2v_i} + 4E_c h_c \frac{1}{r_c}\right) \approx 6850$ Pa, taking $r_c = 8.5$ $\mu$m and $\gamma_m = 40$ pN/$\mu$m [2]. An effective bulk modulus used to model this total elasticity is $K_i = 2283$ Pa. This value is used for the PF simulations in Fig. 4 of the main text, which are quantitatively compared with experimental measurements. However, this $K_i$ requires a very small interface width ($W = 0.16$ $\mu$m) to achieve converged results, which significantly increases computational cost. For most PF simulations, we choose a smaller $K_i = 750$ Pa, as listed in Table S1, and a larger $W$ to improve computational efficiency when studying critical tensions, as they depend only weakly on $K_i$, following a 1/4 power law.

##### E. Numerical implementation

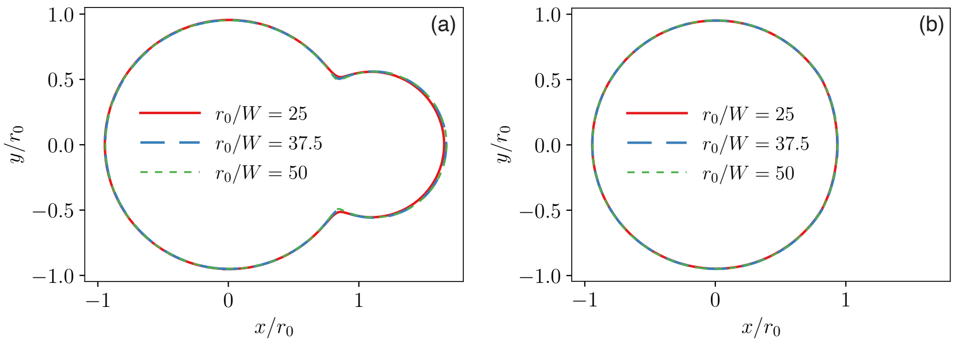

FIG. S2. Comparison of cross-sectional shapes of (a) cell membrane ($\phi_m = 0.5$) and cortex ($\phi_c = 0.5$) in 3D PF simulations with different interface thicknesses for $\lambda = 2$ and $\nu = 12$.

We implement the PF model on massively parallel graphic processing units (GPU) with the computer unified device architecture (CUDA) programming language. The model equations are solved on a square lattice using a finite difference implementation of spatial derivatives and an Euler explicit time stepping scheme. For Laplacian terms in evolution equations, we use isotropic discretizations involving 18 lattice points, including six nearest-neighbor points in $\langle 100\rangle$ directions and twelve next-nearest-neighbor points in $\langle 110\rangle$ directions on a cubic lattice, as described in Ref. [4]. The explicit time step is chosen as $\Delta t/\tau_0 = R_t(\Delta x/W)^2/(12\nu\alpha)$, where the coefficient $R_t$ is set to be 0.5 in all simulations. This choice of $\Delta t$ ensures the numerical stability of Eq. (S10), which has a more stringent stability condition compared with Eq. (S9) for $(\alpha\nu) > 1$.

In PF simulations, we first set both membrane and cortex phase fields to stationary solutions of the form $\phi^0(\vec{r}) = [1 - \tanh(r_0 - |\vec{r} - \vec{r}_{\text{center}}|)]/2$, where $\vec{r}$ is scaled by $W$, $\vec{r}_{\text{center}}$ is the center of the initial spherical cell center, and $r_0$ is the initial cell radius. The initial cell volume $v_0$ is obtained by evaluating the integral $\int d\vec{r} g(\phi_m^0)$. A single perturbed region (suppressed membrane-cortex attachment) is implemented in the PF simulation by setting $\lambda = 0$ in a cylindrical region of radius $r_h$ that extends along the $x$ direction from the cell center to the positive $x$ direction. This creates a circular perturbed region of radius $r_h$ on the periphery of the cell. In simulations with two perturbed regions, we also extend this cylindrical region from the cell center to the negative $x$ direction. To identify the critical tension $T_c$ for endogenous blebbing, we first initiate a bleb by creating a single perturbed region with a small $\nu$. Then we progressively increase $\nu$, which causes a larger $T$, and the bleb can expand beyond the hole region. After the expansion, the bleb solutions are similar with or without perturbation, and we can turn off the perturbation by setting a finite value of $\lambda$ everywhere. Subsequently, we decrease $\nu$ based on the solution of the endogenous bleb and find the critical tension $T_c$ when the bleb spontaneously disappear. For each simulation, we decrease $\nu$ by a small step $\Delta\nu$ (1 or 2). When the bleb solution exists for $\nu_1 = (\nu_0 + \Delta\nu)$ and disappears for $\nu_0$, we measure tensions $T_0$ and $T_1$ corresponding to $\nu_0$ and $\nu_1$, respectively, using Eq. (S21). Thus, we obtain a critical tension $(T_0 + T_1)/2$ and its error $(T_1 - T_0)/2$.

#### II. SHARP-INTERFACE THEORY

In this supplementary section, we develop sharp-interface theories for cellular blebbing. We first discuss the 3D geometry of a cell-bleb system and derive the equation of volume conservation. Next, we examine endogenous blebs, including a sharp-interface analysis of the critical tension required for their formation and a comparison with PF results. Following this, we investigate exogenous blebs, which arise from perturbations in a designated circular region on the cell surface where the membrane-cortex interaction is absent. Finally, we discuss the critical tensions for the formation of such blebs.

##### A. Volume conservation

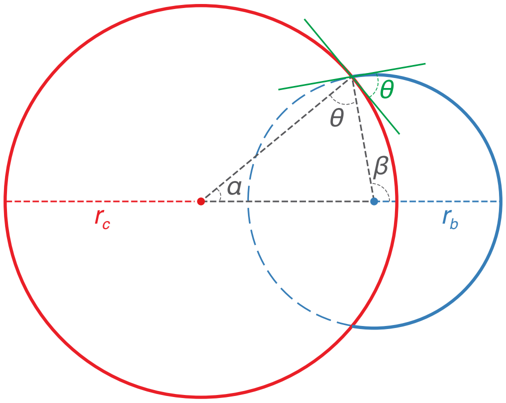

FIG. S3. The cross-sectional view of a 3D bleb, where $r_c$ and $r_b$ are the radii of cell and bleb, respectively, and $\theta$ is the contact angle.

According to the geometrical description of cell and bleb as shown in Fig. S3 (cross-sectional view of a 3D bleb), the overlapping volume of two spheres of radii $r_c$ and $r_b$ is

$$V = \frac{\pi}{12}\frac{(r_c + r_b - d)^2\left(d^2 + 2dr_b - 3r_b^2 + 2dr_c - 3r_c^2 + 6r_br_c\right)}{d}, \tag{S29}$$

where $d$ is the distance between the centers of two spheres. If the $\beta$ angle in Fig. S3 is larger than (or equal to) $\pi/2$, the expression for $d$ is

$$d = \sqrt{r_c^2 - r_h^2} + \sqrt{r_b^2 - r_h^2}, \tag{S30}$$

where $r_h$ is the radius of the circle where two spheres interact. If the $\beta$ angle in Fig. S3 is smaller than (or equal to) $\pi/2$, the expression for $d$ is

$$d = \sqrt{r_c^2 - r_h^2} - \sqrt{r_b^2 - r_h^2}. \tag{S31}$$

Then the bleb volume is calculated according to the change of cell volume:

$$\begin{aligned} V_b =& \frac{4}{3}\pi(r_0^3 - r_c^3) \\ =& \frac{4}{3}\pi r_b^3 - \frac{\pi}{12}\frac{(r_c + r_b - d)^2\left(d^2 + 2dr_b - 3r_b^2 + 2dr_c - 3r_c^2 + 6r_b r_c\right)}{d}, \end{aligned} \tag{S32}$$

where $r_0$ is the initial cell radius. This expression does not explicitly contain the contact angle $\theta$, and it can be calculated according to the relation

$$\theta = \cos^{-1}\left[\frac{(r_c^2 + r_b^2 - d^2)}{2r_c r_b}\right]. \tag{S33}$$

In the limits of $r_b/r_c \to 0$ and $\theta \to \pi/2$, we solve Eq. (S32) for $r_c$ and obtain

$$\frac{r_c}{r_0} \approx 1 - \frac{1}{6}\left(\frac{r_b}{r_0}\right)^3, \tag{S34}$$

which is the same as the solution of $(4/3)\pi(r_0^3 - r_c^3) = (2/3)\pi r_b^3$ that assumes a hemispherical shape of the bleb.

#### B. Endogenous bleb

##### 1. *Equilibrium solutions*

Equilibrium solutions are determined by the force balance condition

$$\frac{2T}{r_c} - K_i\frac{\Delta V}{V_0} = \frac{2\gamma_m}{r_b}, \tag{S35}$$

which follows from equating the pressure in the cell to the Laplace pressure inside the bleb, where $\Delta V = V_0 - V_c = V_b$ with $V_b$ given by Eq. (S32). In the following calculation, we assume $\gamma_m \approx \gamma_m^0$. According to the volume conservation (S34) and the force balance relation (S35), we obtain the relation for a hemisphere bleb ($\theta = 90°$) on a flat interface

$$\widetilde{T} \approx \frac{1}{\tilde{r}_b} + k\frac{\tilde{r}_b^3}{4}, \tag{S36}$$

where $\widetilde{T} \equiv T/\gamma_m^0$, $\tilde{r}_b \equiv r_b/r_0$, and $k \equiv K_i r_0/\gamma_m^0$.

More generally, the relationship between $T$ and $r_b$ (as well as between $T$ and the bleb volume fraction $\rho$) can be determined by numerically solving Eqs. (S32) and (S35). The distance $d$ is determined once we know the contact angle, and for a right contact angle, $d^2 = r_c^2 + r_b^2$. This numerical solution accounts for the full geometry including cortex curvature, and the results are shown as dashed lines in Fig. 2 of the main text.

##### 2. *Critical tensions*

With a sharp-interfacial description, the energy $U$ of the membrane-cortex-cytogel system [Eq. (S5)] reduces to

$$U = TA_{m/c} + \gamma_c A_c + \gamma_m A_m + K_i\frac{(V_0 - V_c)^2}{2V_0}, \tag{S37}$$

where $A_{m/c}$ and $A_c$ are the areas of the membrane- bound and unbound cortex regions, respectively, and $A_m$ is the area of the detached membrane. For a right contact angle, $T = \gamma_c + \gamma_m$, and we obtain

$$U = 4\pi r_c^2 T + 2\pi r_b^2 \left(1 + \frac{d^2 + r_b^2 - r_c^2}{2dr_b}\right)\gamma_m + \frac{K_i}{2}\frac{V_b^2}{4\pi r_0^3/3}. \tag{S38}$$

If we make the approximation that the contact between a hemisphere bleb and the cellular surface can be modeled as a flat interface, which is valid under the condition that $r_b$ is much smaller than the cell radius $r_c$, then the total energy of the system becomes

$$U \approx 4\pi r_c^2 T + 2\pi r_b^2 \gamma_m + \frac{K_i}{6}\frac{\pi r_b^6}{r_0^3}. \tag{S39}$$

According to Eq. (S39), the energy change from the initial value ($4\pi r_0^2 T$) is

$$\Delta U = U - 4\pi r_0^2 T \approx 4\pi T\left(r_c^2 - r_0^2\right) + 2\pi r_b^2 \gamma_m + \frac{K_i}{6}\frac{\pi r_b^6}{r_0^3}. \tag{S40}$$

Assuming $\gamma_m \approx \gamma_m^0$, its dimensionless form is

$$\Delta\widetilde{U} \equiv \frac{\Delta U}{4\pi r_0^2 \gamma_m^0} \approx \widetilde{T}(\tilde{r}_c^2 - 1) + \frac{1}{2}\tilde{r}_b^2 + \frac{1}{24}k\tilde{r}_b^6, \tag{S41}$$

where $\tilde{r}_c \equiv r_c/r_0$. Substituting Eq. (S34) into this equation, we obtain

$$\Delta\widetilde{U} \approx -\frac{1}{3}\widetilde{T}\tilde{r}_b^3 + \frac{1}{2}\tilde{r}_b^2 + \frac{1}{24}k\tilde{r}_b^6 \tag{S42}$$

With the conditions $\frac{d\Delta\widetilde{U}}{d\tilde{r}_b} = 0$ and $\frac{d^2\Delta\widetilde{U}}{d\tilde{r}_b^2} = 0$, we can eliminate $\tilde{r}_b$ and obtain the critical tension for the existence of the endogenous bleb:

$$\widetilde{T}_c^{\text{en}} = \left(\frac{4}{3}\right)^{3/4} k^{1/4}. \tag{S43}$$

*Arbitrary contact angle* If we make the approximation that the contact between a bleb and the cellular surface can be modeled as a flat interface with an arbitrary contact angle $\theta$, then Eq. (S37) becomes

$$\begin{aligned} U \approx & 4\pi r_c^2 T + \pi\left(r_b \sin\theta\right)^2\left(\gamma_c - T\right) + 2\pi r_b^2\left(1-\cos\theta\right)\gamma_m \\ & + \frac{K_i}{2}\frac{[\pi r_b^3(1-\cos\theta)^2(2+\cos\theta)/3]^2}{4\pi r_0^3/3}. \end{aligned} \tag{S44}$$

According to Young's equation $T = (\gamma_c + \gamma_m\cos\theta)$ and $\gamma_m \approx \gamma_m^0$, we can eliminate $\gamma_c$ and obtain the change of the total energy in a dimensionless form:

$$\begin{aligned} \Delta\widetilde{U} \approx & -\frac{1}{6}\left(2 - 3\cos\theta + \cos^3\theta\right)\widetilde{T}\tilde{r}_b^3 - \frac{1}{4}\tilde{r}_b^2 \sin^2\theta\cos\theta \\ & + \frac{1}{2}\tilde{r}_b^2\left(1-\cos\theta\right) + \frac{1}{96}k\tilde{r}_b^6(1-\cos\theta)^4(2+\cos\theta)^2. \end{aligned} \tag{S45}$$

With the conditions $\frac{d\Delta\widetilde{U}}{d\tilde{r}_b} = 0$ and $\frac{d^2\Delta\widetilde{U}}{d\tilde{r}_b^2} = 0$, we can eliminate $\tilde{r}_b$ and obtain the critical tension

$$\widetilde{T}_c^{\text{en}}(\theta) \approx 2\left(\frac{2}{3}\right)^{3/4}\sin\left(\frac{\theta}{2}\right)(2+\cos\theta)^{1/4}k^{1/4}. \tag{S46}$$

As shown in Fig. S4(a), the value of $\widetilde{T}_c^{\text{en}}$ increases with $\theta$ until it stabilizes at a finite value as $\theta \to \pi$. At this point, the bleb is nearly detached from the cell, and $\widetilde{T}_c^{\text{en}}$ becomes independent of $\theta$. In the PF simulation, the interface thickness $W$ influences the critical tension required for blebbing. Fig. S4(a) compares the values of $\widetilde{T}_c^{\text{en}}$ obtained from PF simulations (represented by symbols) for two different values of $W$: 0.213 $\mu$m and 0.32 $\mu$m. For the smaller $W$, the critical tension approaches the sharp-interface predictions corresponding to the saddle-node bifurcation. The sharp-interface predictions include both the analytical solution (S46) for a flat interface (solid line) and the numerical solutions of Eqs. (S32) and (S35) that account for the full geometry (dashed line). The agreement between the latter results and the PF simulations with $W = 0.213\ \mu$m is excellent. Additionally, we plot the scaled bleb radius $r_b/r_0$ as a function of the scaled tension in Fig. S4(b). The PF simulation results for $W = 0.32\ \mu$m (symbols) are compared to the sharp-interface solutions considering full geometry (dotted lines). The agreement between the PF results for blebbing and the sharp-interface solutions worsens as we approach the bifurcation point, particularly for small values of $\lambda$ (i.e., small $\theta$). This suggests that to accurately determine $\widetilde{T}_c^{\rm en}$ in the PF simulations, a sufficiently small value of $W$ is required. Thus, for PF simulations in this study, we use $W = 0.213\ \mu$m unless explicitly stated.

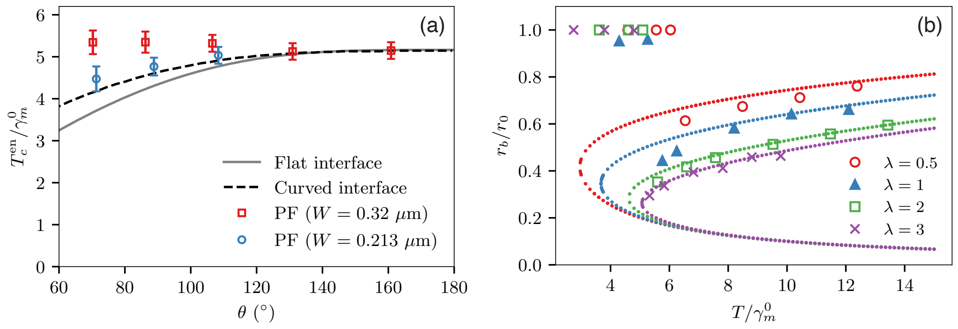

FIG. S4. (a) Critical tension for endogenous blebbing $T_c^{\rm en}$ (scaled by $\gamma_m^0$) as a function of the contact angle $\theta$. Solid and dashed lines are analytical results for a flat and curved interface, respectively, and symbols represent PF results. (b) Scaled bleb radius $r_b/r_0$ as a function of the scaled membrane-cortex tension $T/\gamma_m^0$ for $\lambda = 0.5$, 1, 2, and 3 (measured contact angle $\theta = 40°$, $58°$, $88°$, and $131°$). Symbols represent PF results with $W = 0.32\ \mu$m and dotted lines represent sharp-interface solutions considering the full geometry.

#### C. Exogenous bleb

##### *1. A simple picture*

Next, we investigate the formation of exogenous blebs resulting from permanent perturbations within a specified circular region on the periphery of a cell. In this region, the interaction between the membrane and cortex is absent, mimicking the effects of laser ablation that exogenously ruptures the cortex [2]. For a cell with $N$ perturbed circular regions, each of radius $r_h$, the critical tension $T_c$ is determined when the radius of every bleb, $r_b$ (assuming all $N$ blebs are of identical size), equals $r_h$. Based on volume conservation and the pressure balance relation, the critical tension is given by [2]:

$$\frac{T_c}{\gamma_m} = \frac{r_c}{r_h}\left[1 + N\frac{k}{4}\left(\frac{r_h}{r_0}\right)^4\right] \approx \frac{r_0}{r_h}. \tag{S47}$$

Unlike endogenous blebs, where $T_c^{\rm en}$ is a function of $k$, in this scenario, $T_c$ predicted from this simple picture depends only on $r_h$.

##### *2. $r_h \to 0$ limit*

Here, we consider scenarios where $r_h$ is significantly smaller than both $r_c$ and $r_b$, such that the bleb remains connected to the cell through a circular contact line of vanishing radius, as illustrated in Fig. S5. Under these conditions, the volume conservation equation simplifies to:

$$r_0^3 - r_c^3 = r_b^3. \tag{S48}$$

Since the internal pressures of the cell and the bleb remain equilibrated, we can derive a relation for this type of bleb using Eq. (S35) and Eq. (S48):

$$\widetilde{T} = \frac{1}{2}\left(1 - \tilde{r}_b^3\right)^{1/3}\left(\frac{2}{\tilde{r}_b} + k\tilde{r}_b^3\right). \tag{S49}$$

In the limit $\tilde{r}_b \to 0$, which corresponds to the lower-branch solution, the expression for $\widetilde{T}$ simplifies to:

$$\widetilde{T} \approx \frac{1}{2}\left(\frac{2}{\tilde{r}_b} + k\tilde{r}_b^3\right), \tag{S50}$$

or more simply, $\widetilde{T} \approx 1/\tilde{r}_b$. From Eq. (S49) and the condition $\frac{d\widetilde{T}}{d\tilde{r}_b} = 0$, we obtain the critical tension for the formation of an exogenous bleb in the $r_h \to 0$ limit:

$$\widetilde{T}_c^{\mathrm{ex}} = 2^{1/4}\left(\frac{4}{3}\right)^{3/4} k^{1/4}. \tag{S51}$$

Here, the critical tension is independent of $r_h$ and depends only on $k$.

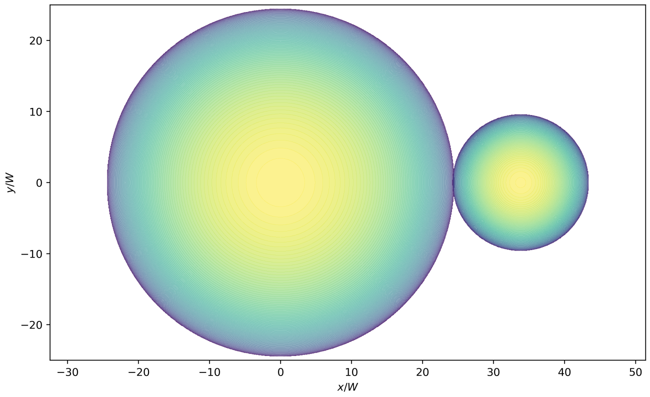

FIG. S5. Blebbing in the $r_h \to 0$ limit.

##### *3. Arbitrary contact angle*

Now, consider a bifurcation picture [Fig. 3(f) in the main text] where the bleb-cortex contact line is anchored at the periphery of a flat perturbed region with a fixed radius $r_h$. Since the primary constraint is the size of the perturbed region, the constraint on the contact angle is relaxed, allowing it to take arbitrary values. For the lower branch solution, i.e., $\theta < \pi/2$, the volume conservation equation [Eq. (S32)] can be approximated as:

$$4\left(r_0^3 - r_c^3\right) = \left(2 - 3c + c^3\right) r_b^3, \tag{S52}$$

where $c \equiv \cos\theta = (1 - r_h/r_b)^{1/2}$. For the middle unstable and upper branches, i.e., $\theta \geq \pi/2$, the volume conservation equation [Eq. (S32)] can be approximated as:

$$4\left(r_0^3 - r_c^3\right) = 4r_b^3 - \left(2 - 3c + c^3\right) r_b^3. \tag{S53}$$

Accordingly, we obtain:

$$\tilde{r}_c = \left[1 - \left(2 + 3bc - bc^3\right)\frac{\tilde{r}_b^3}{4}\right]^{1/3}, \tag{S54}$$

where $b = -1$ for the lower branch and $b = +1$ for the middle unstable and upper branches. By combining the volume conservation condition (S54) with the pressure balance equation (S35), and assuming $\gamma_m \approx \gamma_m^0$ and $r_h/r_0 \ll 1$, we obtain:

$$\widetilde{T} = \frac{\tilde{r}_c}{2}\left[\frac{2}{\tilde{r}_b} - k(\tilde{r}_c^3 - 1)\right]. \tag{S55}$$

Eqs. (S54)-(S55) uniquely determine the bleb radius $\tilde{r}_b$ as a function of tension $\widetilde{T}$, as discussed in the main text.

### III. MULTIPLE BLEBS

#### A. Phase-field simulations of two blebs

In addition to the scenario of a single bleb, multiple blebs on a single cell are also frequently observed in experiments [5]. Here, we investigate the stability of two concurrent endogenous blebs with radii $r_1$ and $r_2$ in contact with a single cell. As shown in Fig. S6(a), we induced two equal blebs on opposite sides of the cell in PF simulations by applying transient perturbations in two circular segments of the cell periphery as the initial condition. The simulation with $\lambda = 2$ demonstrates that the two-bleb system is unstable and eventually reduces to a single larger bleb, which is consistent with the sharp-interface analyzes presented below in Sec. III B 1. Similarly, we examine the stability of two exogenous blebs with $\lambda = 3.5$ under persistent perturbations. Both the PF simulation in Fig. S6(b) and the analysis in Sec. III B 2 indicate that this configuration is unstable. The only stable two-bleb configuration corresponds to the solution in the lower branch, which is accessible only under persistent perturbations. This result is confirmed by both the sharp-interface analysis in Sec. III B 2 and the PF simulation in Fig. S6(c), where the blebs in the lower branch of solutions are significantly smaller.

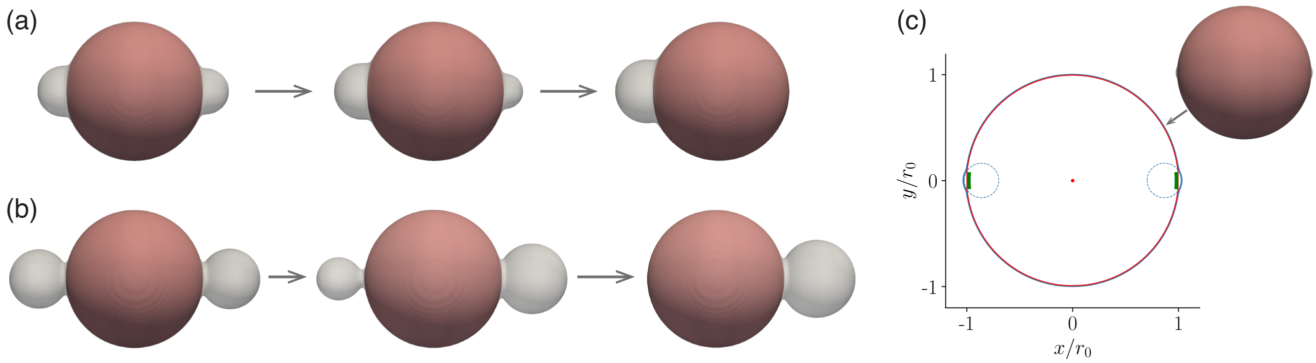

FIG. S6. PF simulations of the two-bleb systems. (a) The elimination process of one of the endogenous blebs, as observed in a PF simulation with $\lambda = 2$ and $\nu = 8$. (b) The elimination process of one of the exogenous blebs, as observed in a PF simulation with $\lambda = 3.5$ and $\nu = 14$ under persistent perturbations, i.e., suppressing membrane-cortex attachment in two small circular regions of radius $r_h/r_0 = 0.08$. (c) The lower stable branch of solution in a PF simulation with $\lambda = 3.5$ and $\nu = 8$. The region with suppressed membrane-cortex binding is marked as a green patch ($r_h/r_0 = 0.08$) in the cross-sectional view from the 3D PF simulation.

#### B. Stability of two-bleb configurations

##### 1. *Endogenous blebs*

We investigate the stability of a two-bleb configuration where each endogenous bleb belongs to the upper stable branch of solutions in Figs. 2(a)-(b) of the main text. If we make the approximation that their interaction with the cellular surface can be modeled as a flat interface, which is valid under the condition that the radii $r_1$ and $r_2$ of both blebs are much smaller than the cell radius, $r_c$. Conservation of volume implies $(4/3)\pi(r_0^3 - r_c^3) = (2/3)\pi(r_1^3 + r_2^3)$, which yields

$$r_c \approx r_0 \left(1 - \frac{1}{6}\left[\left(\frac{r_1}{r_0}\right)^3 + \left(\frac{r_2}{r_0}\right)^3\right]\right), \tag{S56}$$

in the limit $r_i \ll r_0$ $(i = 1, 2)$. For this two-bleb system, the energy change from the initial state is

$$\Delta U \approx 4\pi T(r_c^2 - r_0^2) + 2\pi\gamma_m(r_1^2 + r_2^2) + \frac{K_i}{6}\frac{\pi(r_1^3 + r_2^3)^2}{r_0^3}. \tag{S57}$$

Combining Eqs. (S56) and (S57), we eliminate $r_c$ and obtain the change of total energy in its dimensionless form:

$$\Delta\widetilde{U} \equiv \frac{\Delta U}{4\pi r_0^2\gamma_m^0} \approx -\frac{1}{3}\widetilde{T}(\tilde{r}_1^3 + \tilde{r}_2^3) + \frac{1}{2}(\tilde{r}_1^2 + \tilde{r}_2^2) + \frac{1}{24}k(\tilde{r}_1^3 + \tilde{r}_2^3)^2, \tag{S58}$$

where the approximation $\gamma_m \approx \gamma_m^0$ is used. The equilibrium point of this two-bleb system in the $(\tilde{r}_1, \tilde{r}_2)$ space is found when $\tilde{r}_1 = \tilde{r}_2 = \tilde{r}$ and $\Delta\widetilde{U} = 0$, which yields

$$-4\tilde{r}\widetilde{T} + 6 + k\tilde{r}^4 = 0. \tag{S59}$$

The stability at the equilibrium point is determined by the determinant of the Hessian matrix, which is given by

$$H = \begin{bmatrix} \frac{\partial^2 \Delta\widetilde{U}}{\partial \tilde{r}_1^2} & \frac{\partial^2 \Delta\widetilde{U}}{\partial \tilde{r}_1 \partial \tilde{r}_2} \\ \frac{\partial^2 \Delta\widetilde{U}}{\partial \tilde{r}_2 \partial \tilde{r}_1} & \frac{\partial^2 \Delta\widetilde{U}}{\partial \tilde{r}_2^2} \end{bmatrix}. \tag{S60}$$

For $k = 150$, the numerical calculation for different $\widetilde{T}$ shows that the determinant of $H$ is negative. This indicates that the system has a saddle point at the equilibrium point, indicating that the system is unstable. The contour plot of $\Delta\widetilde{U}$ is shown in Fig. S7. This prediction agrees well with the PF simulation in Fig. S6(a).

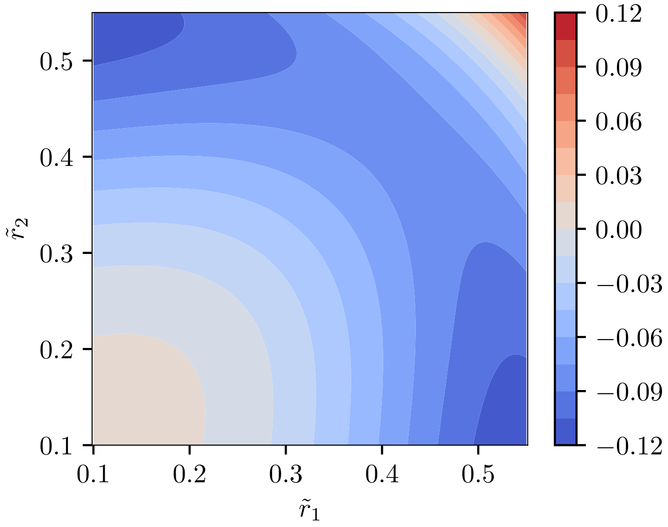

FIG. S7. Contour plot for the change of the total energy $\Delta\widetilde{U}$ in a two-bleb system with $k = 150$ and $\widetilde{T} = 8$. The axes represent the scaled radii of each bleb, denoted as $\tilde{r}_1$ and $\tilde{r}_2$.

##### *2. Exogenous blebs*

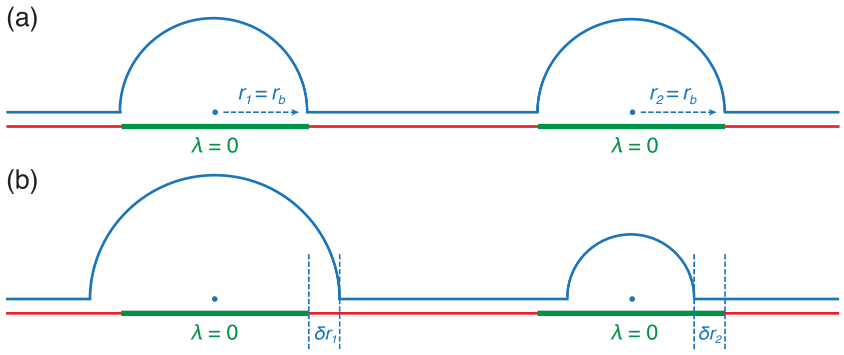

FIG. S8. Illustration of a two-bleb system with two perturbed regions: (a) the initial state where the radii of two blebs ($r_1 = r_2 = r_b$) are equal; (b) two blebs with unequal sizes.

*a. Lower stable branch of solutions.* We first consider a two-bleb configuration where each small bleb belongs to the lower stable branch of solutions in Figs. 3(a)-(b) of the main text and the tension is just below $T_c$. In this case, the two blebs can be assumed to be hemispherical with the same initial radii $r_1 = r_2 = r_h$. A perturbation analysis is performed based on this state with a positive variation in the radius of the first bleb, $\delta r_1 > 0$ as shown schematically in Fig. S8. Assuming the variation is small, volume conservation ensures that the corresponding variation in the radius of the other bleb satisfies $\delta r_2 \approx -\delta r_1$ at linear order. In addition, we consider for simplicity the case where $T \approx \gamma_c$ where $\theta = \pi/2$ (e.g., $\lambda = 2$ in Fig. 1c). With $\lambda = 0$ in the perturbed region, the combined tension of the membrane and cortex is simply $\gamma_c + \gamma_m$ (since there is no binding energy), which exceeds $T$. Thus, the change in total energy at linear order is

$$\delta U \approx 2\pi r_h \gamma_m \delta r_1 > 0, \tag{S61}$$

which implies that this two-bleb configuration is energetically stable. This prediction agrees qualitatively well with the PF simulation in Fig. S6(c), which shows that a configuration of two exogenous blebs from the lower stable branch of solutions is stable with a finite $r_h$.

*b. Upper stable branch of solutions.* Next, we consider a two-bleb configuration where each large bleb belongs to the upper stable branch of solutions in Figs. 3(a)-(b) of the main text. This case is simpler to analyze in the $r_1 = r_2 = r_h \to 0$ limit. A perturbation analysis is performed as before based on this state with a positive variation in the radius of the first bleb, $\delta r_1 > 0$. The volume conservation ensures that the corresponding variation in the radius of the other bleb satisfies $\delta r_2 = -(r_1/r_2)^2 \delta r_1$. In the $r_h \to 0$ limit, only the change of the bleb surface area accounts for the variation of the total energy of this two-bleb system. Thus, the change in total energy is

$$\delta U = -8\pi\gamma_m \left(\frac{r_1}{r_2} - 1\right) r_1 \delta r_1 < 0, \tag{S62}$$

which implies that this two-bleb configuration is energetically unstable. This prediction in the $r_h \to 0$ limit agrees qualitatively well with the PF simulation in Fig. S6(b), which shows that a configuration of two exogenous blebs from the upper stable branch of solutions is unstable even when $r_h$ is finite.

*
